# Anatomical and Palynological Investigation within flora of Family *Asteraceae*

**DOI:** 10.1101/2022.04.22.489184

**Authors:** Asim Shahzad, Shahid Iqbal, Sadaf Kayani, Tayyab Shafiq, Muhammad Zafar, Muhammad Naeem, Humaira Yasmin

## Abstract

A comprehensive microscopic investigation of the leaf epidermis, as well as anatomical and palynological research of selected *Asteraceae* species from the flora of Havali (Kahutta) Azad Jammu and Kashmir (AJK) Pakistan, was carried out. This study includes 11 plant species comprising of *Senecio jacobea, Leucanthemum vulgare, Halianthus annuus, Erigeron bonariensis, Achillea millifolium, Halianthus Linnaeus, Taraxacum officinale, Anaphalius nepalensis, Erigeron Canadensis*, and *Tagetes erecta*. All the species studied were amphistomatic, with four different forms of stomata i.e tetracytic, anomocytic, anisocytic, and tricytic. The main stomatal type was tetracytic, followed by anomocytic. The abaxial epidermis has a higher stomatal density than the adaxial epidermis. Highest stomatal density was present *Tagetes erecta* while lowest is present in *Conyza canadensis*. In lower epidermis stomatal index was higher in *Taraxacum officinale* followed by *Halianthus annuus* and *Tagetes erecta* while in upper epidermis highest index was shown by *Halianthus annuus*. For this study species of family *Acteraceae* were properly collected. Furthermore, variation in pollen can be seen. List of palynomorph which includes family name, botanical name, local name, English name, flower colour, season and pollen description for the logical arrangement of these species. The logical ordering of these species was guided by dust characteristics such as form and pollen morphology. Palynological data has been found to be too big for taxonomists to calculate and make appropriate observations on their findings.

## 1. Introduction

*Asteraceae* is a large family with over 1500 genera and 25,000 species in various parts of the world [1]. Spices, subshrubs, and bushes, as well as trees, are all members of this family [2]. Epidermal characteristics of leaves, such as somatic cell dividers, stoma types, and trichomes, are key boundaries at both traditional and explicit levels in diverse plant groups. Stomata are holes formed by a few unique cells called watchman cells that are situated on the surface of aeronautical sections of upper plants and may open and close gas exchange between the plant and its environment. Their purpose is to act as entry points for CO_2_ into the leaf for photosynthesis and exit points for water fumes, which might be used for evaporative cooling. The flowing water may also aid in the uptake and delivery of salts required for the plant’s survival [3].

Stomatal circulations are influenced by the status of the plant’s surroundings. The leaves of amphibian-lowered plants do not have stomata. Stomata opens when photosynthesis is happening at a good rate and closes when there isn’t enough water. Trichrome is a kind of epidermis that is glandular in nature. This might be constant and present in all parts of the plant’s body. Trichomes might possibly be used in the identification of plant species [4]. There is a variation in stomatal size due to the size of guard cells. Stomata are holes that may be found on both the upper and lower epidermis and are bordered by a pair of guard cells. The major purpose of epidermal pores is to exchange gases like carbon dioxide and oxygen between plants and the environment [5]. Amphistomatic leaves with hexacytic stomata were discovered in the basal epidermis of family Polyonaceae [6]. In dicots, there are seven types of stomata, with the amphianisocytic form being the most frequently observed [7]. The density of ordinary epidermal cells and stomata in the epidermis of a shadow leaf was lower than that of a leaf present in sunlight. Furthermore, stomatal thickness was positively related to cell length and stomata holes [8].

Palynology is simply the study of the accumulation of “thrown particles.” All things considered, an exceptional palynologist evaluates particle models derived from the air, water, or deposits, including residue. The state and identity of typical inorganic “thrown particles” provide palynologists with clues about the life, condition, and lively conditions that transported them.

The first known declaration of residue under an amplifying focus is thought to be in the 1640s by English botanists [9]. Pollen is essential for sexual dissemination in flowering plants, according to those who depicted residue and therefore the stamen and assumed that pollen place sampling. Specialists have mentioned a variety of common tree dust kinds, as well as many spores and taste residual grains. A study of residue tests obtained from the accumulation of Swedish lakes is being conducted [10], they discovered so much pine and clean residue that he believed they may be used as “record fossils.”

Light and filtering microscopy were used to investigate the dust morphology of seven species from the greenery of Kaghan valley that shared a neighborhood with seven *Asteraceae* genera [10]. *Linn, Chrysanthemum leucanthemum, Linn, Gerbera gossypina (Royle) Beauv, Senecio chrysanthemoides DC, yarrow Linn, Chrysanthemum leucanthemum, Linn, Gerbera gossypina (Royle) Beauv, Senecio chrysanthemoides, Sonchus asper Linn., African marigold Linn* and *Xanthium strumarium Linn* [10]. The simplest dust size was discovered in G. gossypina, which was 38.3 m in polar view and 57.75 m in central view, while the smallest dust size was found in *X. strumarium*, which was 23.65 m in polar view and 19 m in tropical view during a *Millefolium* study [11].

The objectives of the present investigation were to investigate the Study of the pollen variation of family *Asteraceae* in Havali (Kahutta) and to study the micro-morphological interaction and degree of variations in the leaf epidermis of preferred species of *Asteraceae* and the basis of leaf epidermal characteristics a step to pre-flowering identification.

## 2. Materials and Methods

The Forward Kahutta (33.8844° N, 74.1034° E) which is the hot spot of this study is the administrative unit (sub-divisions) of the District Haveli. In addition, numerous locations (villages) were chosen for survey and data gathering including Lasdana, Badori, Haji Peer, and Aliabad.

### 2.1. Collection of study materials

Field data was collected between January and December 2019 during visits to the research region and through interviews with the locals. The legal authority to conduct the interview in this research was obtained from “district and local government authorities.” The “prior informed consent” (PIC) documents were translated into Havali’s regional tongue.

### 2.2. Isolation of epidermis

To prevent dehydration, young leaves were submerged in water. The abaxial and adaxial epidermis of each exemplar were isolated using a razor, needles, and forceps [6]. In several species, a coat of direct nail cleaner was truly effective on the leaf surfaces. Within the period of drying, the concept revealed an astonishing aspect of skin [12].

### 2.3. Staining, mounting and microscopic examination

Leaf tests were done by using the protocols present in previous literature with slight modifications [13]. New leaves were used and after drying they were placed in high temperature water by using water shower until become fragile and opened. Both dry and fresh leaves were percolated for 2–3 minutes in a tube stacked with 10% nitric destructive and 90% lactic acid, till the models were clear. Camel brush was used to carefully remove the adaxial and abaxial surfaces. The epidermis was placed at the glass slide and observed under the light microscope for the immediate evaluation. For each model, five occurrences of the two surfaces were discovered, and its different attributes were observed. The images were taken by employing camera (Meiji CCD Model 00179048, Canada) fitted on Leica Dialux 20 light amplifying instrument. By and large, trichome expressed itself [14]. Three readings of each component were noted for the 2 surfaces. This gives information about the length and width of the foliar epidermal features.

### 2.4. Method of studying palynology

#### 2.4.1. Collection, identification, and preservation

Surveys were conducted to rare wetlands in District Havali Azad Jammu and Kashmir, where humans can influence the *Asteraceae* family in the changing seasons. Wetlands at Badori, Lasdana, Haji Peer, and Aliabad have been visited for the variety of flora. Sprouting mannequin instances grouped together with the help of their natural surroundings. After grouping, the flowers are dried and crushed with the help of smirching sheets. Plant species were identified using distinguishing and herbarium analyses from Pakistan’s Herbarium (ISL). Flora of Pakistan also confirmed the existence of the species [15]. Plants are pressed, dried, and connected to herbarium sheets before being submitted to the Herbarium of Pakistan (ISL) at Quaid e Azam University in Islamabad, Pakistan (Table 1).

**Table 1:** Floristic Composition and Trichome Type of Selected Species of Asteraceae.

| S.no | Plant Species | English name | Flower<br>color | Season of<br>flower | Locality<br>Altitude |
| --- | --- | --- | --- | --- | --- |
| 1 | *Senecio jacobaea* | Tansy ragwort | Yellow | June and<br>July | Bedori-3727 m |
| 2 | *Leucanthemum*<br>*vulgare* | Bellis perennis | White | Early summer<br>mid summer | Lasdana-2625m |
| 3 | *Helianthus annuus* | Sun flower | Yellow | Summer | Haji Peer -3421m |
| 4 | *Erigeron bonariensis* | Argentine Fleabane | Blue, cream &<br>purple | Summer | Bedori-3727 |
| 5 | *Achillea millefolium* | Yarrow | White | May to June | Alibad-2682m |
| 6 | *Helianthus Linnaeus* | Sun Flower | Yellow or<br>Mahroon | Summer | Lasdana-2625 |
| 7 | *Taraxacum officinal* | Bitterwort | Yellow | Summer to<br>Spring | Haji Peer -3421m |
| 8 | *Anaphalius*<br>*nepalensis* | Spreng | White | Summer | Alibad-2682m |
| 9 | *Erigeron Canadensis* | Canadian fleabane | Cream,<br>Yellow | Jully to<br>October | Badori-3727 |
| 10 | *Tegetes erecta* | African Marigold | Yellow | Summer | Lasdana-2625 |

#### 2.4.2. Pollen analysis through light microscopy

Residue grains from floral blossoms are splattered on a glass slide and a drop of acid was poured. With the help of glycerin jam, a slide was once mixed and recolored. The residue grains are counted and analyzed by Leica Dialux 20. The unusual features like polar, focal separation, exine thickness, length, and breadth of colpi, and holes length were observed after the examination of tidies. The tidies’ position was determined by the distance between polar and tropical separation. Size and situation of the residue grains has been taken with the help of (Punt, Hoen, Blackmore, Nilsson, and Le Thomas, 2007). SPSS writing PC packages offered a wide range of related recommendations to observe and eradicate general mistake to examine the records.

### 2.5. Data analysis

The data was tabulated and analyzed using the formulae listed below, and all statistical findings were evaluated with statistix 8.1.

#### 2.5.1. Determination of stomatal complex types

Stomatal complex was studied on the abaxial and adaxial epidermis of the selected species [16].

#### 2.5.2. Determination of stomatal density and index (%)

“The number of stomata per square millimeter of leaf surface” is stomatal density.

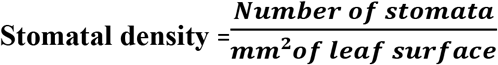

“Number of stomata divided by number of epidermal cells multiplied by 100” is stomatal index (%) [17].

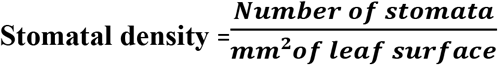

## 3. Results

A total of ten plants belonging to the *Asteraceae* family were examined for Palynomorph features in this study. Palynomorph stock is found in alphabetical scales of families, followed by characteristic name, English name, Palynomorph highlights the botanical name, English name, floral colour, and season of 10 species in table 1. Ten *Asteraceae* species were investigated for microscopic features of leaf epidermis.

### 3.1. Microscopic analysis of Stomatal features of selected species of *Asteraceae*

The amphistomatic species were those with the same or separate stomatal types on the abaxial and adaxial epidermis. Stomatal edifices of four types were studied: tetracytic, anomocytic, anisocytic, and tricytic. Tetracytic stomata were the most common, followed by anomocytic, anisocytic, and tricytic stomata (Table 2). Stomatal thickness was larger on abaxial epidermis than adaxial epidermis in numerous species (Figure.3). *H. annuus* (199.16) had the highest stomatal thickness, followed by *T. erecta* (195.01) and *E. bonariensis* (111.09), respectively. *H. annuus* (201.17) had the highest stomatal thickness on the adaxial surface, followed by *T erecta* (132.94) and *E. bonariensis* (115.08).

**Figure 1.**
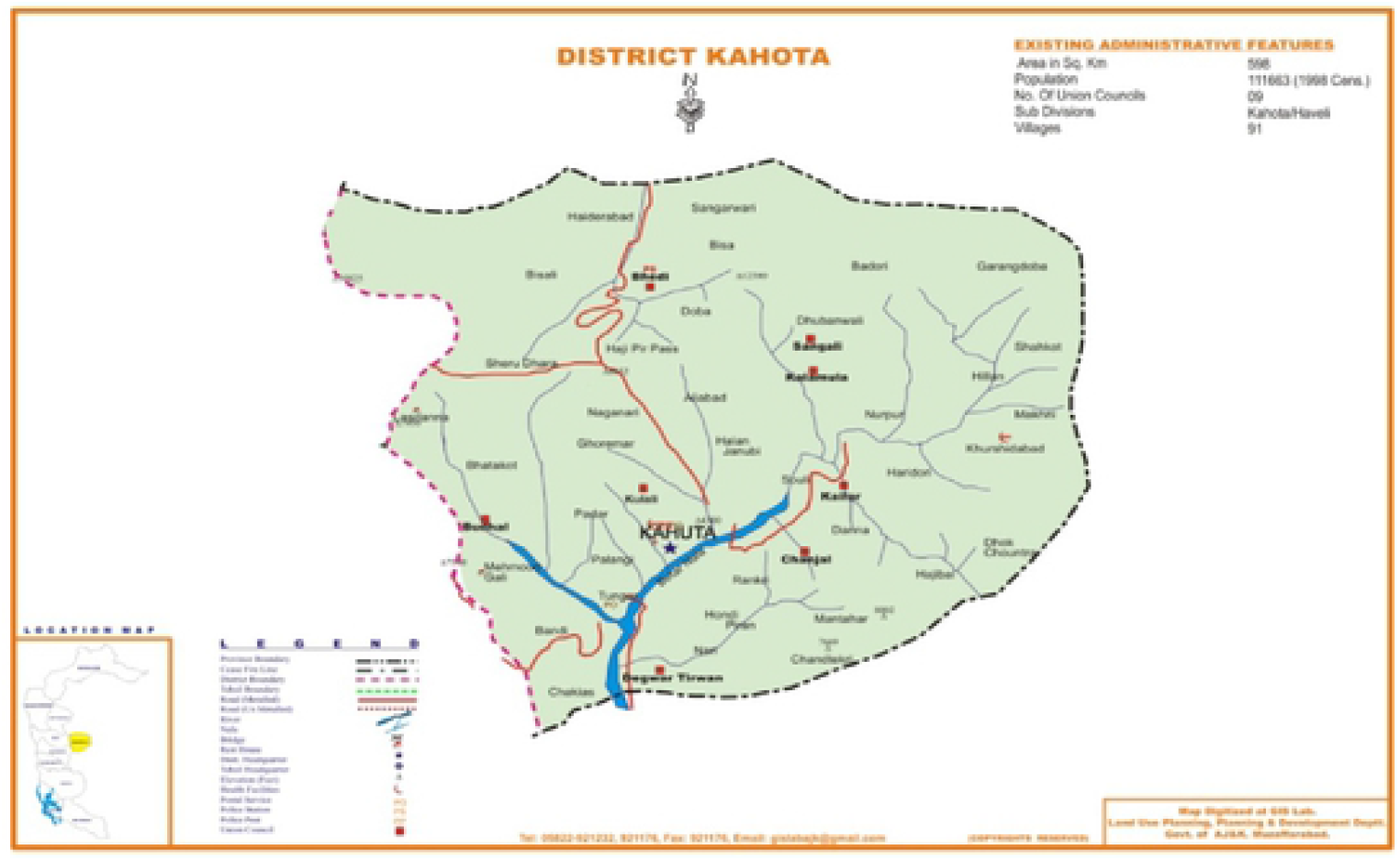
Map of Havali (Kahutta) Azad Kashmir.

**Figure 2.**
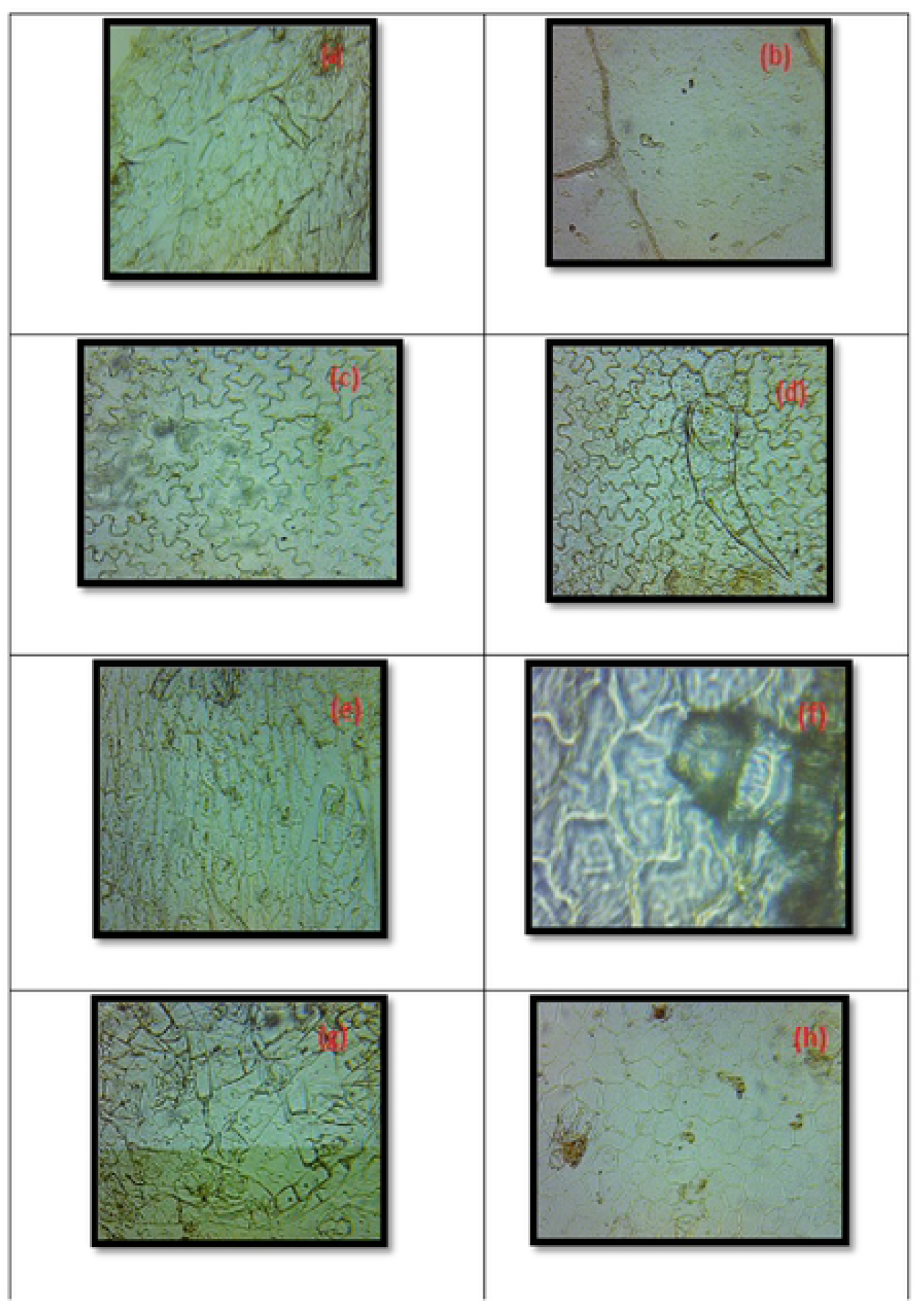
Abaxial and Adaxial view of epidermal anatomical of selected species. a. *Senecio jacohaea* b. *Leucanthemum vulgare* c. *Halianthus annuus* d. *Erigeron honariensise. Achillea millefolium* f. *Halianthus Linnaeus* g. *Taraxacum officinale* h. *Anaphalius nepalensis* i. *Erigeron Canadensis* j. *Tegetes erecta*

**Figure 3.**
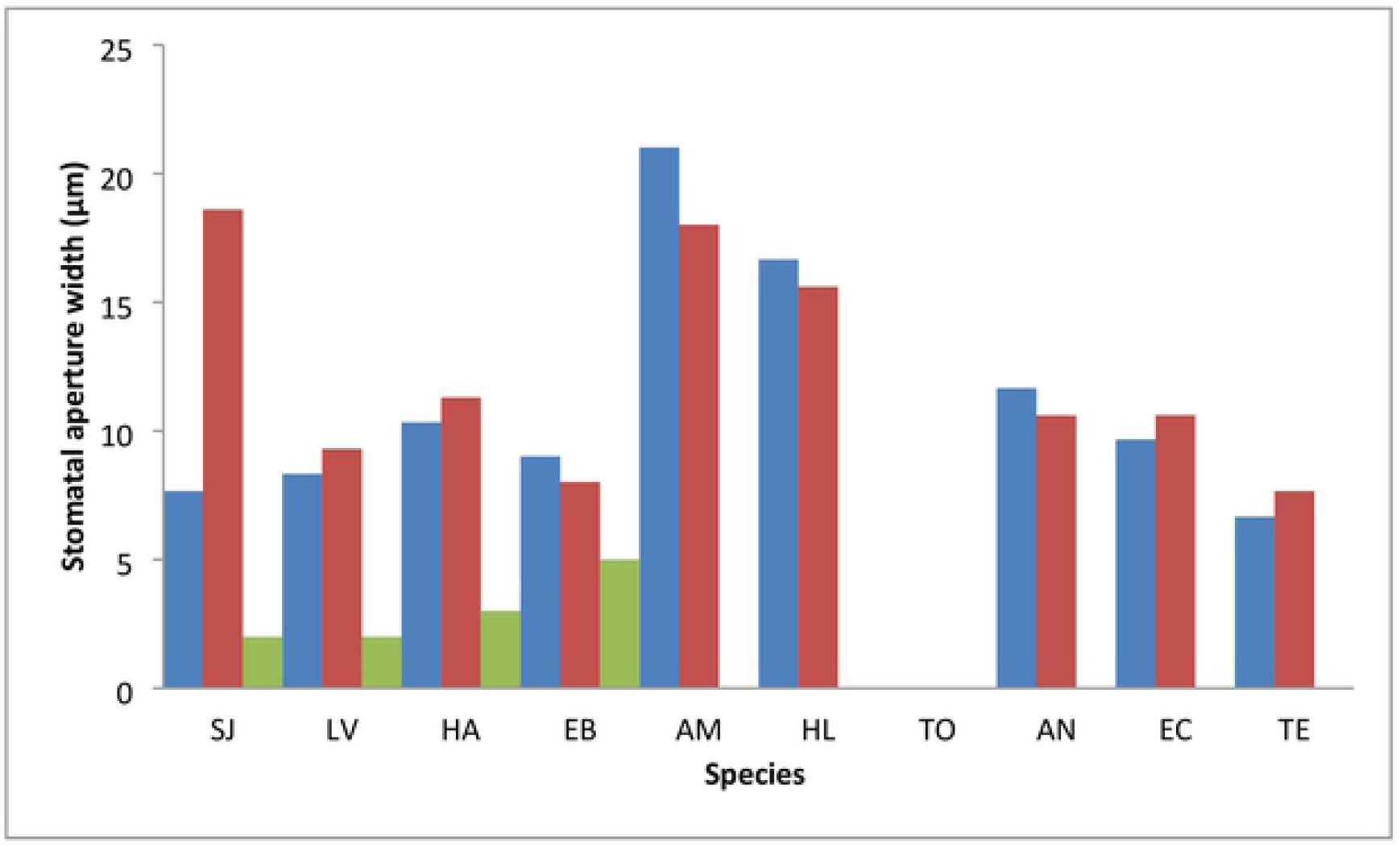
Stomatal aperture width of selected Asteraceae species. SJ= Senecio jacohaea, LV= Leucanthemum vulgare, HΛ= Halianthus annuus, EB= Erigeron bonariensis, AM= Achillea millefolium, HL= Halianthus Linnaeus, TO= Taraxacum officinal, AN= Anaphalius nepalensis, EC= Erigeron Canadensis, TE= Tegetes erecta

**Figure 4.**
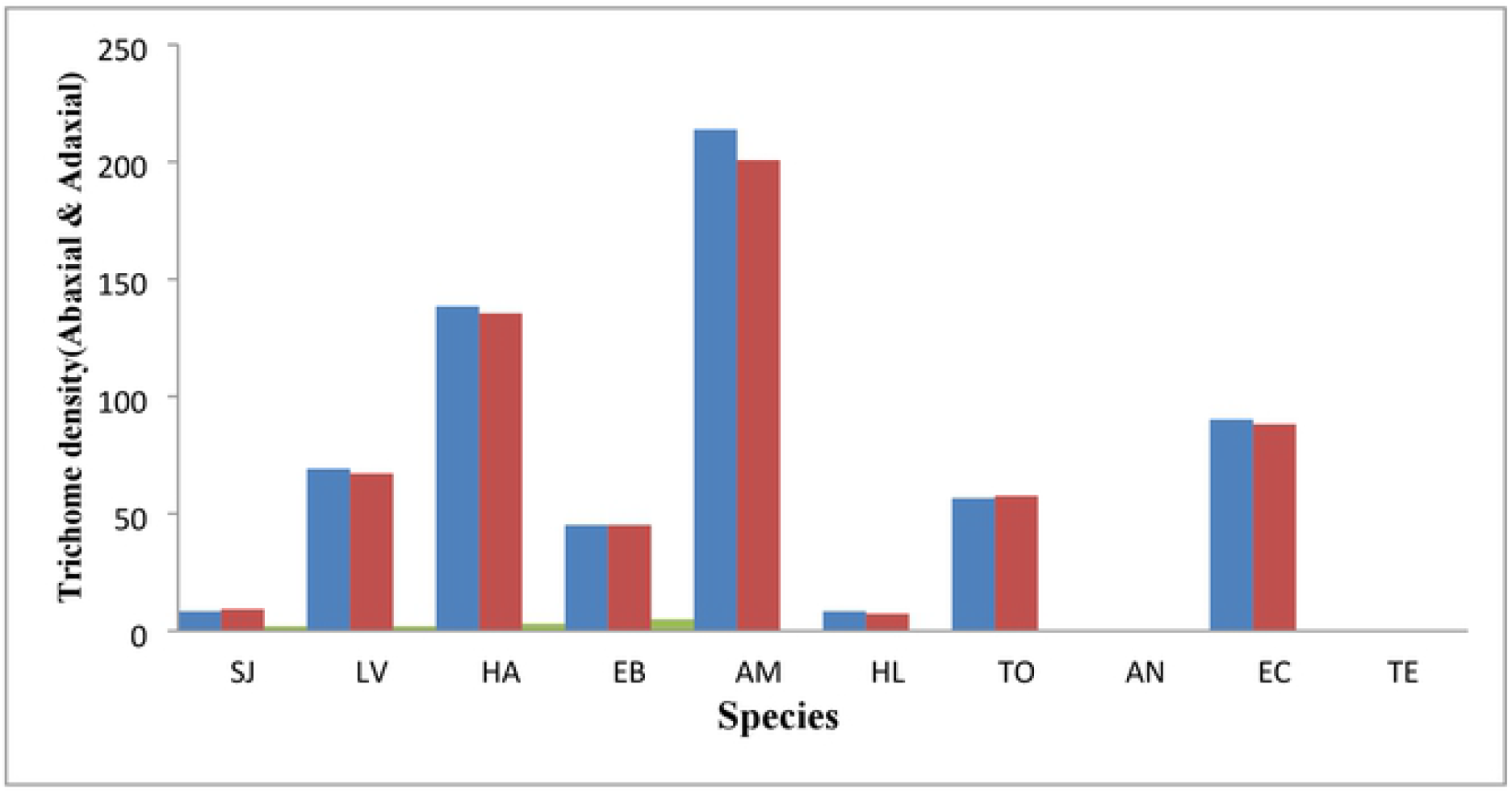
Trichome density of selected Asteraceae species. SJ= Senecio jacohaea, LV= Leucanthemum vulgare, HA= Ilalianthus annuus, EB= Erigeron bonariensis, AM= Achillea millefolium, HL= Ilalianthus Linnaeus, TO= Taraxacum officinal, AN= Anaphalius nepalensis, EC= Erigeron Canadensis, TE= Tegetes erecta.

**Figure 5.**
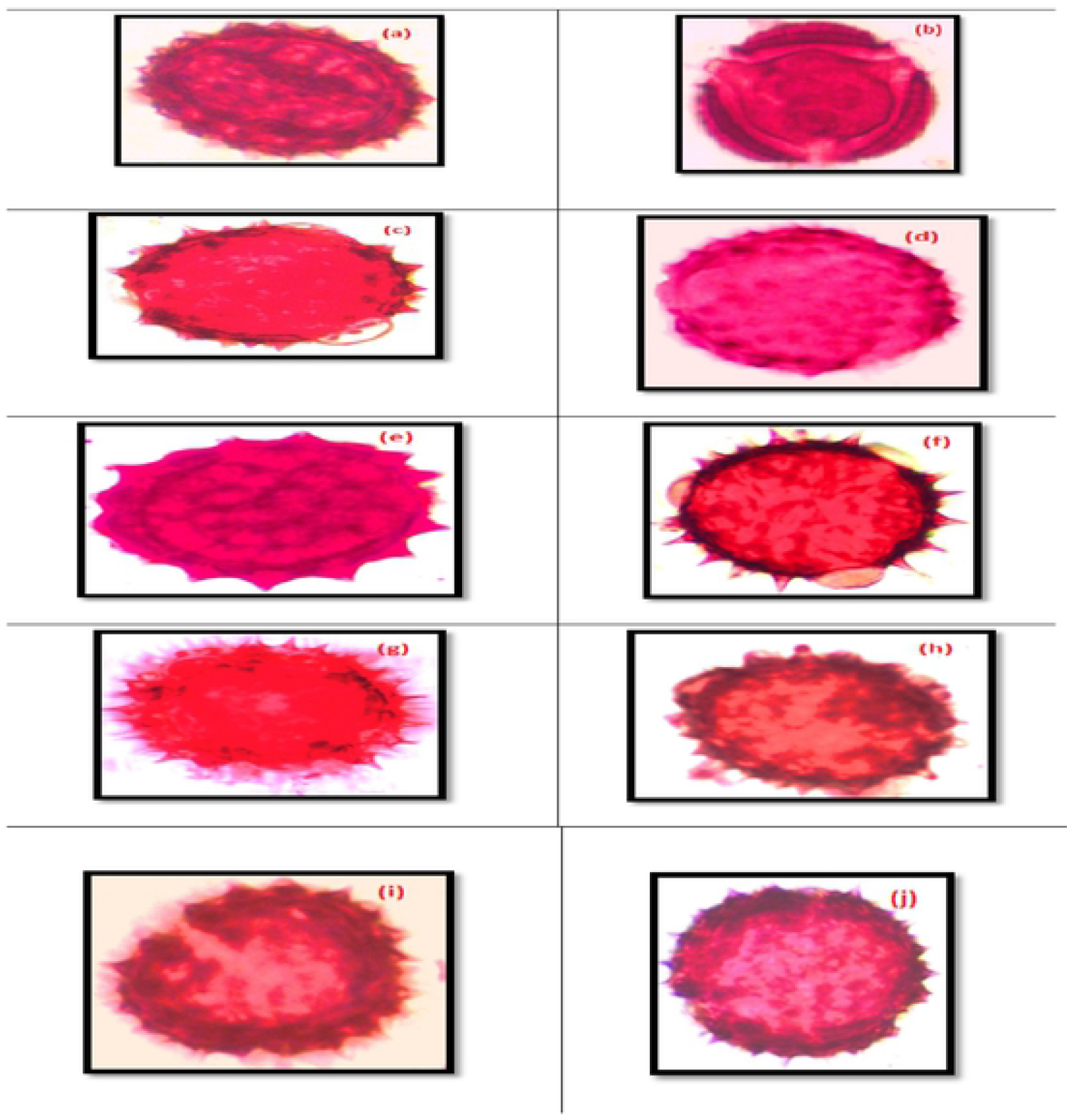
Polar and equatorial view of pollen selected species. a. *Senecio jacohaea* b. *Leucanthemum vulgarec. Halianthus annuus* d. *Erigeron honariensise. Achillea millefolium* f. *Halianthus Linnaeus* g. *Taraxacum officinale* h. *Anaphalius nepalensis. Erigeron Canadensis* j. *Tegetes erecta*

**Table 2:**
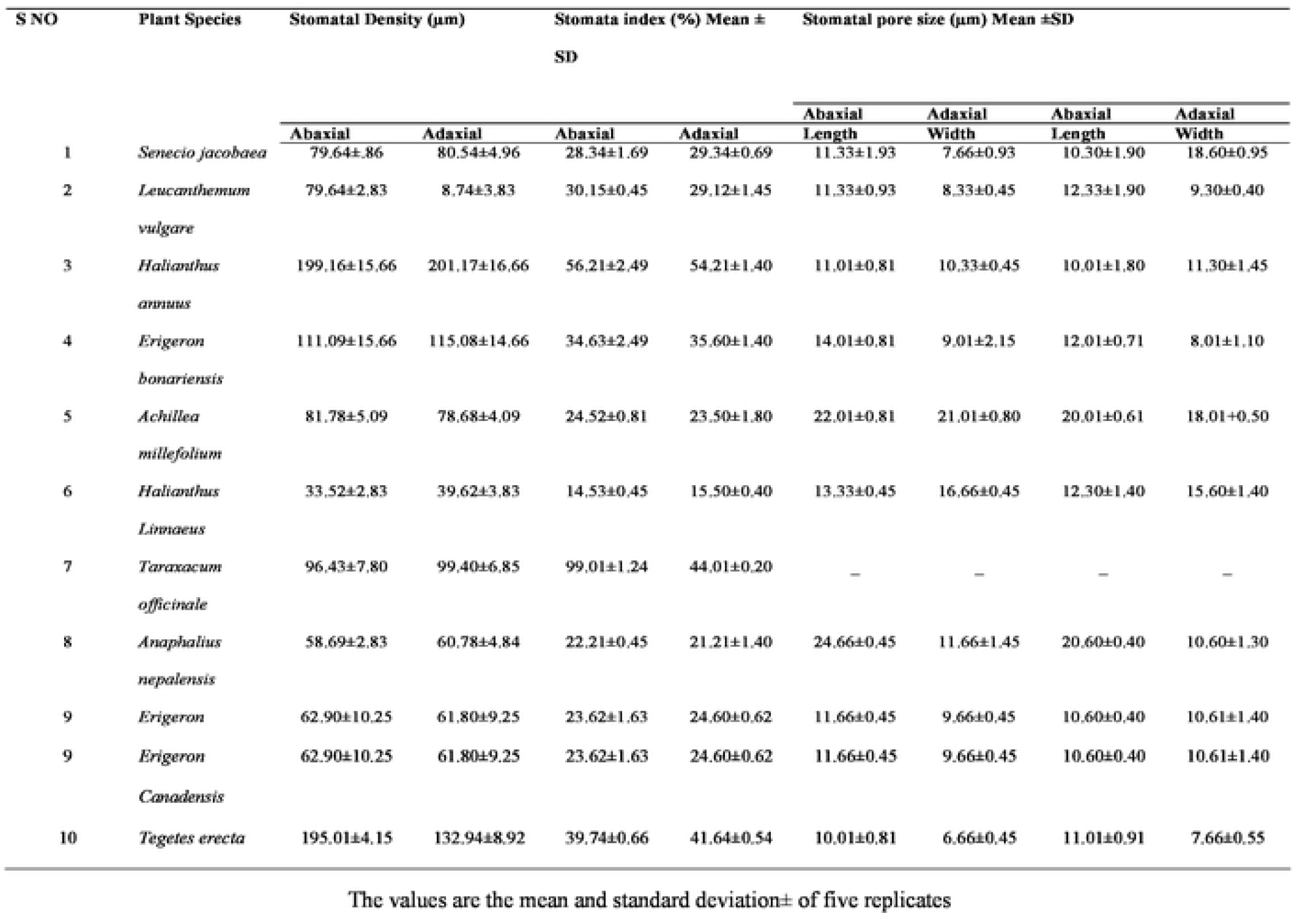
Microscopic analysis of Stomatal features of selected species of Asteraceae.

On the abaxial epidermis, *H. linnaeus* (33.52) had the smallest stomatal thickness, whereas on the adaxial epidermis, *H. linnaeus* (39.62) had the most. In the lower epidermis, *T. officinal* (99.01 *%*) was the highest, followed by *H. annuus* (56.21 *%*) and *T. erecta* (39.74 *%*), while in the upper epidermis, *H. annuus* (54.21 *%*), *T. oppicinal* (44.01 *%*), and *T. erecta* (41.64 *%*) were the highest seen in table 2. On the abaxial epidermis, the best pore was found in *A. nepalensis* (24.66 m), followed by yarrow *A. millefolium* (22.01 m) and *E. bonariensis* (14.01 m), while the pore length was shorter in *T. erecta* (10.01 m). In the abaxial epidermis of *A. millefolium* (21.01 m), *H. linnaeus* (16.66 m), and *A. nepalensis* (11.66 m), more expanded stomatal pores were seen, whereas in the adaxial epidermis of ragwort (18.60 m), and of *A. millefolium* is (18.01 m).

### 3.2. Microscopic Analysis of Trichome of Selected Species of *Asteraceae*

Four distinct trichomes were discovered. Multicellular with acicular culmination (Senecio jacobaea), multicellar with top pointed oxeye daisy, yarrow, and horseweed, multicellular summit globular *E. bonariensis, H. Linnaeus*, and unicellular turn (*H. annuus, T. officinal*). Multicellular with acute apex, multicellular with spherical apex, multicellular with acicular, and unicellular spine are the trichome types listed in table 3. *A. millefolium* (213.88) had the thickest trichomes on the abaxial epidermis, followed by *H. annuus* (138.39) and *E. Canadensis* (90.14). *S. jacobaea*, on the other hand, has the smallest population (8.36). *H. annuus* (135.39) had the highest trichome thickness on the adaxial epidermis, followed by *A. millefolium* (130.80) and *E. Canadensis* (88.14) seen in Figure 2.

**Table 3.** Trichome Type of Selected Species Asteraceae.

| S.no | Plant Species | Trichome type |  |
| --- | --- | --- | --- |
|  |  | Abaxial | Adaxial |
| 1 | *Senecio jacobaea* | Unicellular, Spinee like | Unicellular, Spinee like |
| 2 | *Leucanthemum vulgare* | Multicellular, Apex globular | Multicellular, Apex globular |
| 3 | *Helianthus annuus* | Multicellular, Apex pointed | Multicellular, Apex pointed |
| 4 | *Erigeron bonariensis* | Unicellular, Apex globular | Unicellular, Apex globular |
| 5 | *Achillea millefolium* | Unicellular, Spine like | Unicellular, Spine like |
| 6 | *Helianthus Linnaeus* | Absent | Absent |
| 7 | *Taraxacum officinal* | Multicellular, Apex pointed | Multicellular, Apex pointed |
| 8 | *Anaphalius nepalensis* | Absent | Absent |
| 9 | *Erigeron Canadensis* | Unicellular, Spinee like | Unicellular, Spinee like |
| 10 | *Tegetes erecta* | Multicellular, Apex globular | Multicellular, Apex globular |

### 3.3. Microscopic analysis of Pollen features of selected species of *Asteraceae*

In a wide range of *Asteraceae* species, dust is almost spherical and spheroidal. The immensity in headway and at an unmistakable and ordinary level arranged by this family are the characteristics of residue spine. As shown in table 4, spineless residue was detected in *L. vulgare* and *T. officinale*. In polar perspective, the most extravagant residue size, for example, 12.3 m, was discovered in *T. officinal*, while the smallest residue size, for example, 7.6 m, was found in *A. nepalensis*, as indicated in table 4. Most extreme dust size for example 12.1 μm was found in *H. Linnaeus* in tropical view and least dust size for example 6.8 μm in equatorial view was watched in *E. Bonariensis* and *E. Canadensis* as appeared in table 4. Maximum ratio i.e., 1.17 in *F. bonariensis* and *E. candansis* and minimum is 0.88 in *H. Linnaeus* shown in Figure 2.

**Table 4.**
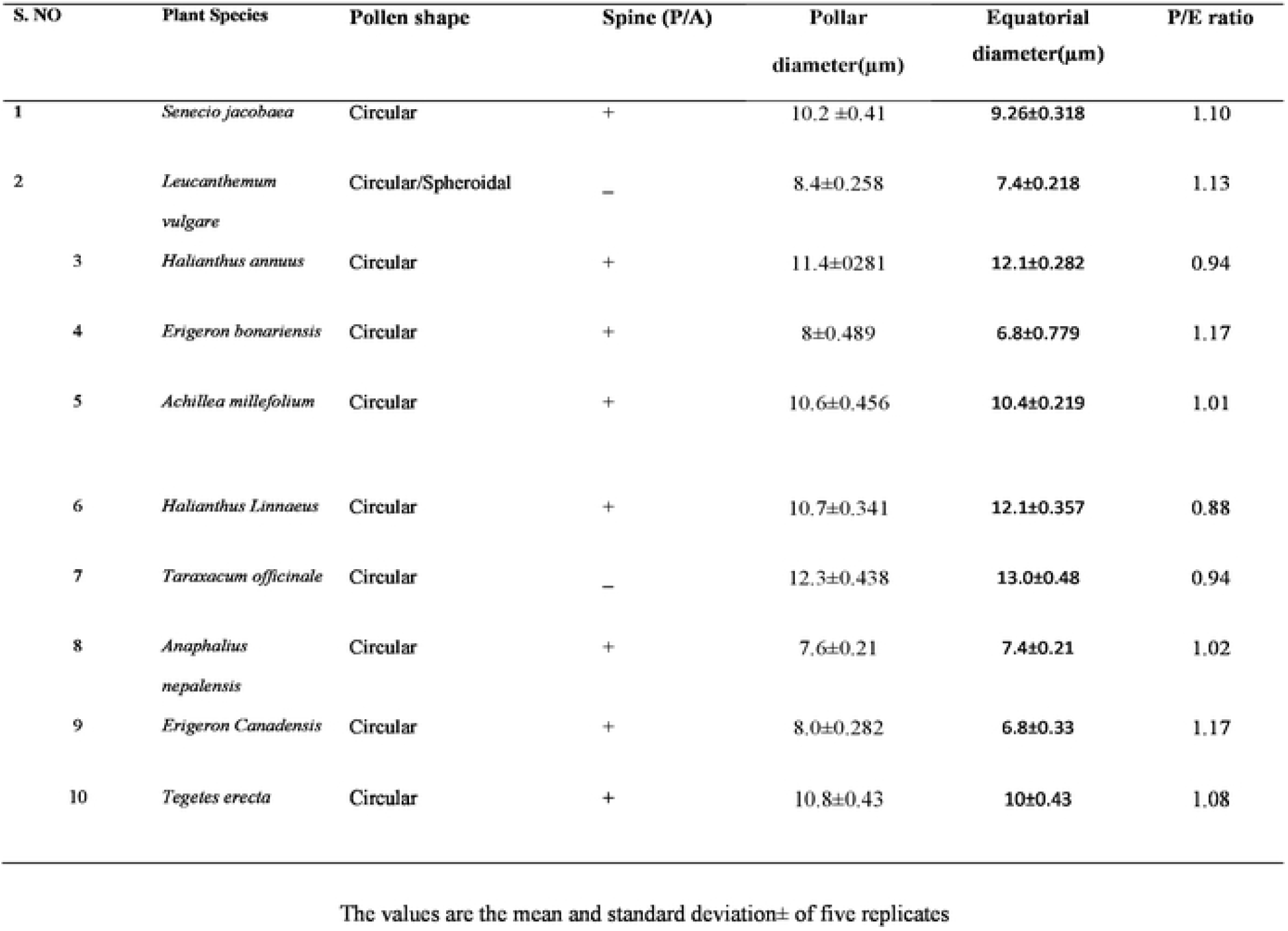
Microscopic analysis of Pollen features of selected species of Asteraceae.

### 3.2. Microscopic analysis of Colpi and spine feature of selected species of

Number of colpi in all selected species is same and number of spines is shown in table 5. In *H. Linnaeus* biggest spine length was seen which is 2.9 μm was and typically negligible in *E. bonariensis* and *A. nepalensis* which was 0.6 μm, similarly the foremost vital colpi length was 2.9 μm in *A. millefolium*, whereas humblest in colpi length was found in *T. officinale* that was 4.75 μm. showed up in table 5. Maximum distance between two colpi is 8.4 μm in *H. Linnaeus* and min is 5.2 μm in *E. bonariensis* shown in table 5. Max no of spine between colpi is 9.2 μm in *T. ereacta* and min is 4.4 μm in *A. millefolium* shown in table 5.

**Table 5.** Microscopic analysis of Colpi and spine feature of selected species of Asteraceae.

| S No | Plant Species | No of colpi | No of spine | Spine length (μm) | Colpi length (μm) | Distance between two colpi (μm) | Number of spin between clopi |
| --- | --- | --- | --- | --- | --- | --- | --- |
| 1 | *Senecio jacobaea* | 3 | 26.2±1.7 | 0.88±0.076 | 2.3±0.21 | 5.44±0.60 | 8.4±0.21 |
| 2 | *Leucanthemum vulgare* | 3 | — | — | 1.2±0.204 | 5.8±0.437 | — |
| 3 | *Halianthus annuus* | 3 | 24±1.019 | 0.78±0.037 | 1.1±0.139 | 5.74±0.548 | 6±0.282 |
| 4 | *Erigeron bonariensis* | 3 | 24.2±1.706 | 0.6±0.039 | 1.5±0.130 | 5.2±0.334 | 6.6±0.357 |
| 5 | *Achillea millefolium* | 3 | 18.6±0.606 | 1.14±0.20 | 2.9±0.089 | 6.6±0.357 | 4.4±0.219 |
| 6 | *Halianthus Linnaeus* | 3 | 31.2±1.729 | 2.9±0.0071 | 1.2±0.036 | 8.4±0.456 | 8±0.2817 |
| 7 | *Taraxacum officinale* | 3 | 30.32±1.682 | 1.18±0.132 | 1.6±0.150 | 7.3±0.57 | 8.2±0.331 |
| 8 | *Anaphalius nepalensis* | 3 | 22±0.66 | 0.62±0.10 | 1.1±0.05 | 5.4±0.2 | 5.6±0.21 |
| 9 | *Erigeron Canadensis* | 3 | 21.6±1.45 | 0.74±0.035 | 1.0±0.08 | 5.8±0.43 | 6.6±0.21 |
| 10 | *Tegetes erecta* | 3 | 30.4±0.96 | 0.9±0.12 | 2.5±1.87 | 7.2±0.52 | 9.2±0.43 |

The values are the mean and standard deviation± of five replicates

## 4. Discussion

The stomatal life structures of the leaf epidermis were combined to measure the degree of assortments in selected types of Asteraceae constructed from region Havali AJ&K. Stomata are found on both surfaces of the leaf in these species, with irregular, uneven, parallel, and cross cells [18]. All of the selected species had at least one distinct stomata near the mirasol, in line with previous research on the *Ficus* sp. [19]. Stomatal opening length changed from 10.01 μm to 24.12 μm and 10.30 μm 20.01 μm on abaxial and adaxial epidermis, uninhibitedly these discernment appear as if to the finding of [20]. On the dorsal and ventral epidermis, the width of the aperture decreased from 7.66 m to 21.01 m and from 7.66 m to 18.66 m, respectively, autonomously concurring with the presentations in *Polygonaceae* as provided by [6]. Stomatal length on the dorsal and ventral epidermis displays anatomical similarities between the leaf surfaces. The upper stomatal record was found in the lower epidermis of T. *ruderalia* and in the upper epidermis of H. *annus*. Stomatal degrees from 14.53% to 99.01% in selected species, It was analyzed that stomatal index of some species of this family higher on lower epidermis and on upper epidermis of other species as reported in Onasma kind of *Baranginaceae* [21]. Multicellular trichomes are referenced further into three sorts enthusiastic to cell shapes situated at on apex, these cell of multicellular trichomes are sharp and spherical. *Asteraceae* species have sharp, spherical and with one, pair, many cell trichome [18].

On abaxial and adaxial epidermis of comparable species assortment were viewed. Within most of the species stomatal features were consistent. Basic differences showed up within the stomatal complex sorts, thickness, record, and size of pore. Trichome thickness is missing in *A. nepalensis* and *T. erecta*. Also, stomatal pore size missing in *T. officinale*. These arrangements are useful to disengage these species.

Palynology is particularly related with the appraisal of astute event, paleobotany, ethnobotany, inherited and upgrades studies, arranging and climatic changes and condition. Dust morphology is a technique with which developing grains are frequently created for examination. Movement and identification of vegetations from diverse places and parties are highlighted by taxonomists [22]. In selected types of *Asteraceae*, the development shape is mostly deviant and spheroidal. This family’s characteristic of development spine is criticalness, which is formed and ordered at an unmistakable and usual level. Oxeye daisy and common dandelion both showed signs of development. The spinate dust character is taken under consideration as a foul fragment when stood separated from yellow development. The current assessment produced a record that was confined to unambiguous palynological evaluations of Compositae development morphology. Taxonomists from any location who examine the greeneries of a clear district of their inclinations would benefit from such research. The *Asteraceae* family is eurypalynous, and most of its taxa contain zonocolporatepollen. This research discovered a surprising intentional variety in *Asteraceae* dust morphology, including differences in size, shape, spine length, quantity, and colpi morphology. Development size as an example 12.3 μm was found in *T. officinal* in polar view and least development size as an example 7. 6 μm in polar view was seen in *A. nepalensis*. Despite the way that the bulk phenomenal development size in focal view is 13 μm in *T. officinal* and least was 6.8 μm in both *E. bonariensis* and *horseweed*.

Most conspicuous spine length was seen in *H. Linnaeus*, which was 2.9 μm and each one around diminished in *E. bonariensis* and *A/ nepalensis*, which was 0.6 μm, additionally the foremost essential colpi length as an example 2.9 μm in yarrow, whereas humblest in colpi length was found in *E. Canadensisthat* was 1.0 μm, whereas in *P. hysterophours* the colpi length wasn’t without a doubt seen. It’s captivating to need a gander at the connection of long spines with high colpi length. If this alliance exists it’d be of much arranged criticalness. Rest of the species exhibited midway examinations of spine and colpi length. Our conceded consequences of spine plan are according to the finding of [23] and [12], who used spine length and number of spine pushes between colpi as observing referenced character in palynology of Compositae. Spines within the type Calendula are basically name, i. e. a couple of enduringly orchestrated spines are accessible during this class [23]. Such spines are out and out illustrative to determine this tax structure other taxa during a specific gathering. Such assessments have become a massive, designed contraption due to sensible variation in dust morphology.

### Conclusion

There are significant differences between the stomatal complex types, density index, size of pore and trichomes of ten selected species of *Acteraceae* namely *S. jacobaea, L. vulgare, H. annuus*, *E. bonariensis*, *A. millefolium*, *H. Linnaeus*, *T. officinale*, *A. nepalensis, E. Canadensis*, *T. erecta*. To differentiate these species these variations are very helpful. Most of the characteristic are steady in same class and in equivalent species variation were observed on abaxial and adaxial epidermis. In some species of *Asteraceae*, the shape of trichome is important character.

## Conflicts of interest/Competing interests

Authors declare that no conflict of interest exists

## Ethics approval

Not Applicable

## Consent to participate

All authors consent to participate in this manuscript

## Consent for publication

All authors consent to publish this manuscript in the Saudi Journal of Biological Science

## Availability of data and material

Data will be available on request to the corresponding or first author

## Code availability

Not Applicable

## Authors’ contributions

**AS=** writing—original draft, methodology, data curation, formal analysis, conceptualization; **SI**= review and editing, conceptualization, supervision, project administration; **TS**= Writing original draft, review and editing, statistical analysis, **SK**=administration, review and editing, **MZ**= formal analysis, review and editing, resources, **MN**= review and editing, statistical analysis **HY**= Resources, data curation, formal analysis, review and editing

## Acknowledgments

The findings of this study are a part of the MS studies of above-mentioned students.

## Notes

### Competing Interest Statement

The authors have declared no competing interest.

